# Functional disruption of Transferrin expression alters reproductive physiology in *Anopheles culicifacies*

**DOI:** 10.1101/2021.06.14.448311

**Authors:** Jyoti Rani, Tanwee Das De, Charu Chauhan, Seena Kumari, Punita Sharma, Sanjay Tevatiya, Soumyananda Chakraborti, Kailash C Pandey, Namita Singh, Rajnikant Dixit

**Affiliations:** Laboratory of Host-Parasite Interaction Studies, ICMR-National Institute of Malaria Research, Dwarka, New Delhi, 110077, India; Department of Biotechnology, Guru Jambheshwar University of Science & Technology, Hisar, India

**Keywords:** Mosquito, Blood-Meal, Iron-metabolism, Immunity, Reproductive Physiology

## Abstract

Iron metabolism is crucial to maintain optimal physiological homeostasis of every organism and any alteration of the iron concentration (i.e. deficit or excess) can have adverse consequences. Transferrins are glycoprotein’s that play important role in iron transportation and have been widely characterized in vertebrates, and insects, but poorly studied in blood-feeding mosquitoes. Here, we characterized a 2102 bp long transcript *AcTrf1a* encoding putative transferrin homolog protein from mosquito *An. culicifacies*. A detailed *in silico* analysis predicts *AcTrf1a* (ACUA023913-RA) encodes 624 amino acid (aa) long polypeptide that carries transferrin domain. *AcTrf1a* also showed a putative N-linked glycosylation site, a characteristic feature of most of the mammalian transferrin’s and certain non-blood feeding insects. Structure modelling prediction confers the presence of an iron binding site at the N-terminal lobe of the transferrin. Our spatial and temporal expression analysis under altered pathophysiological conditions showed that *AcTrf1a* abundantly express in the fat-body, ovary, and its response is significantly altered (enhanced) after blood meal uptake, and exogenous bacterial challenge. Additionally, a non-heme iron supplementation of FeCl_3_ at 1 mM concentration not only augmented the *AcTrf1a* transcript expression in fat-body, also enhanced the reproductive fecundity of gravid adult female mosquitoes. RNAi mediated knockdown of *AcTrf1a* causes a significant reduction in the egg laying/fecundity, confirmed important role of transferrin in oocyte maturation. Further detailed characterization may help to select this transcript as a unique target to impair the mosquito reproductive outcome.

**Highlights:**

1. Insect transferrins are mostly glycoprotein of about 60-80 kDa molecular weight, involved in myriad physiological events and serve as a major iron transport protein.
2. Here, we identified and characterized a 2102 bp long transcript encoding putative transferrin homolog of 624 aa long peptide, carrying only one fully functional transferrin domain at N-terminal from *An. culicifacies*.
3. Spatial and temporal expression analysis of *AcTrf1a* highlights an enriched expression in fat-body and ovary during vitellogenesis.
4. Iron supplementation and dsRNA mediated knockdown experiments together confer that *AcTrf1a* may have key role in the iron homeostasis regulation during oogenesis, and egg maturation in the gravid female mosquitoes.

**Graphical abstract:** Fig 1:
Schematic presentation of iron transport from midgut to ovary by transferrin1 and oocyte reduction after *AcTrf1a* knockdown.
Mosquito acquires iron either from blood meal or iron supplementation in sugar meal. Fat-body derived transferrin proceed towards the gut surface, load iron in its N-terminal iron-binding pocket and deliver iron to ovary. This blood meal iron is required by adult female for completion of gonotrophic cycle. (a) limited iron availability in sugar meal does not support the ovary development and hence no oogenesis; (b) when sugar meal is replaced by blood meal upregulation of transferrin protein results in rapid iron transport to various organs including ovary results in healthy ovarian growth; (c) RNAi mediated knockdown of this transporter protein transferrin in fat-body followed by blood meal, may cause reduced iron transport to ovary and consequently declines in oocyte load.

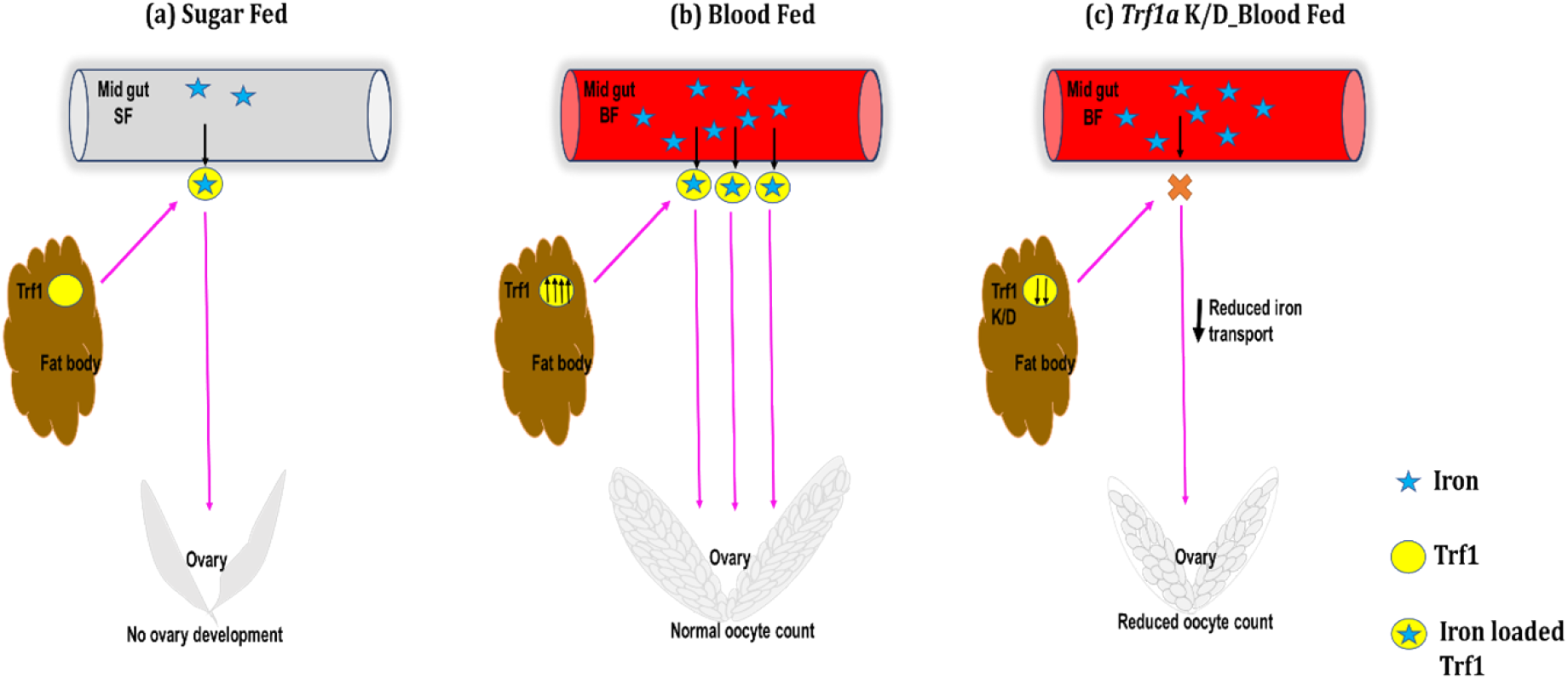

## Introduction

Iron is a trace biologically indispensable and potentially toxic element. All living organisms require iron for innumerable biological functions such as energy generation *via* electron transport, oxygen transport, DNA synthesis and repair, immunity, healing, cuticle formation, melanization, and as a biocatalyst for various processes (1–3). In addition, in anautogenous insects, blood meal iron is an essential requirement for the ovarian follicles growth and oocyte development (4, 5). A surplus amount of iron leads to hydroxyl radical production which may cause cell/tissue damage, and impact overall performance of an individual. Therefore, maintaining an optimal level of iron is crucial for survival of every biological organism (6–9). Several studies on vertebrate system highlight that transferrin is a major iron transport carrier in the blood (10, 11). Among the four well characterized mammalian transferrins (serum transferrin, lactotransferrin, melanotransferrin and inhibitor of carbonic anhydrase) only transferrin and lactotransferrin are secretary in nature, and binds to iron with a greater affinity (12). Although, transferrin superfamily members have been reported in more than 34 invertebrate species, including insects, very little is known about its role in iron transportation in mosquito species (13, 14). Among all those protein involved in iron transport Transferrin and Ferritin gained most attention, because of their unique structure and their unique iron binding ability (1).

Insect transferrins, having molecular weight of ∼ 66 kDa, are identified as juvenile hormone regulated protein, and function as potent immune protein, antibiotic agent, antioxidant, vitellogenic protein and provide protection from plant secondary metabolite etc., (15, 16). *Kurama et. al*., have suggested that transferrin is involved in vitellogenesis in flesh flies (*Sarcophaga peregrina*), utilized by developing oocytes (17, 18). Beside these roles, transferrin protein is also found as utmost component of milk composition in tsetse flies for intrauterine larval development (19). However, unlike vertebrate transferrin, insect transferrin evolved with only N- terminal iron-binding pocket for iron-binding and transportation (20, 21). Although this strategy, seems to sacrifice half of the iron-binding capacity of the transferrin to defeat the “piracy” action of the pathogen, but molecular basis of iron transportation is still sparse (22).

Anautogenous mosquitoes take iron-rich blood meal for their egg maturation and completion of gonotrophic cycle (23–25). Rapid utilization of this heavy blood meal iron content may also result in the formation of hydroxyl free radicals that are highly destructive (26, 27). To minimize this oxidative effect insects are evolved with a unique ability to mobilize, utilize and store iron through various iron-binding proteins, such as ferritin (storage) and transferrin (transportation) (8). Insect homolog transferrin has also been identified from several mosquito species, and their immune role has been explored in few mosquitoes such as *Ae. aegypti* (28, 29); *An. gambiae*, and *Cu. quinquefasciatus* (30). For e.g., an *in-vitro* study on *Cu. pipiens pallens*, showed prolific expression in cypermethrin resistance strain, advocating that transferrin may confer insecticide resistance against cypermethrin (31). However, so far functional correlation of transferrin in the reproductive physiology of any mosquito species remains unexplored. Literature suggests that in many insects (including *D. melanogaster, Ae. aegypti* and *An. gambiae*) transferrin 2, 3 and 4 are co-existing with transferrin1 (15).

*An. culicifacies*, is a major vector responsible for 65% malarial cases in rural India (32). Control of *An. culicifacies* is very challenge because of development of insecticide resistance, and biology of this mosquito remains poorly understood. In the present study, we describe *An. culicifacies* transferrin 1 (*AcTrf1a)* which shares the highest 80% identity with *An. stephensi* followed by 74%, 52% and 21% identity with *An. gambiae, Ae. aegypti* and *Ho. sapiens* transferrin, respectively (Supplementary table 6). We also reported the differential expression of *AcTrf1a* transcript in response to development, blood meal, iron and bacterial challenge. Finally, using functional genomics approach we showed that *AcTrf1a* play important role in shaping the reproductive fecundity of mosquito *An. culicifacies*.

## Material methods

### *In silico* analysis: Domain arrangement, localization and structural modelling of *AcTrf1a*

The putative *AcTrf1a* gene was retrieved from the hemocyte RNA-Seq data of naïve *An. culicifacies*. Initial BLASTx analysis against the NCBI NR database showed a significant hit to the transferrin-like proteins originating from multiple mosquitoes and insect species. To retrieve full-length transcript, BLASTn analysis was performed against the genome predicted transcript database of the *An. culicifacies* mosquito, which is available on *www.vectorbase.org*. Top 10-15 blast hits FASTA sequences were selected from mosquito and non-mosquito species, followed by alignment using ClustalX2 for multiple sequence alignment analysis. The phylogenetic tree was generated by providing aligned file of ClustalX2 as an input in maximal likelihood programme of MEGAX software. The programme was run on 1000 boot straps to confers branching reliability (33). *AcTrf1a* domain annotation was performed using online web server Pfam (http://pfam.xfam.org/). Iron-binding site of *AcTrf1a* was determined using sequence alignment with human transferrin (GenBank ID: NP_001054.1). CASTp (http://sts.bioe.uic.edu/castp/index.html?1bxw) was used for protein cavity analysis. Homology modelling of *AcTrf1a* (sequence and other details of *AcTrf1a* was provided in Supplementary Table 1 and 2) was performed using three independent automated online server Phyre2 (http://www.sbg.bio.ic.ac.uk/~phyre2/html/page.cgi?id=index), RaptorX (http://raptorx.uchicago.edu/) and Robetta (https://robetta.bakerlab.org/). The quality of different protein models developed by different servers was assessed by SAVES v6.0 (https://saves.mbi.ucla.edu/) and Robetta generated model was found the best and selected for further analysis. Structural visualization of protein was carried out using Pymol (https://pymol.org/2/). N-linked glycosylation site prediction was done by Gene Runner software on default parameters and further validated by CBS prediction servers (http://www.cbs.dtu.dk/services/TargetP/).

### Mosquito rearing

*An. culicifacies* (sibling species A) was reared and maintained in central insectary of NIMR under standard rearing condition of 28 ± 2°C, relative humidity 60-80% and 12:12 hr light/dark cycle. Aquatic stages were reared in enamel trays with water supplemented with mixture of dog food and fish food. After emergence adults were kept in cages and fed on cotton swab dipped in 10% sugar solution. Ethical committee of NIMR has approved all the steps taken for rearing and maintaining cyclic colony of mosquitoes.

### Bacterial challenge assays

Exogenous bacterial challenge was posed by injecting bacterial culture of *Escherichia coli* and *Bacillus subtilis* in 0-2 days old female mosquitoes using nanoinjector facility (Drummond Scientific Nanoject II, Broomall, PA, USA). Both *E. coli* and *B. subtilis* were cultured in Luria Bertani broth as overnight culture at 37°C. Bacterial culture at log phase were centrifuged to 20000g for 10min. After washing pellet 2-3 times with phosphate buffered saline (PBS) these were resuspended in PBS buffer. For heat-killed assays bacterial cells dissolved in PBS having OD-0.91 and 0.96 for *E. coli* and *B. subtilis*, respectively, were kept at 90° −120°C for 10 min to expose pathogen-associated molecular patterns (PAMPs). Mosquitoes after challenge with heat-killed bacterial soup, were placed in plastic cups, having moistened surface with wet filter paper-pad, and covered with net. Wet cotton swab and raisins were kept on the net for better recovery under standard rearing conditions. The experimental design consisted of two different treatment groups: the ‘bacterial challenge’ group was injected with 69 nl of bacterial solution and the control group with 69 nl of PBS buffer from same cohort of mosquitoes. Post injection, around 30-35 mosquitoes were dissected for hemocyte tissue collection in time dependent manner. Each group and time point were replicated three times.

### Exogenous Ferric chloride feeding assay

For non-heme iron feeding, mosquitoes just after emergence were either fed on 10% sucrose solution alone (control) or on sugar supplemented with FeCl_3_ till the end of the experiment. Initially three different concentrations of ferric chloride – 1 μM, 5 μM and 10 μM were selected to check the responsiveness of the selected transcript in its origin tissue. But no significant modulation was observed. Afterwards, all experiments were done using the 0.1 mM, 0.5 mM and 1 mM conc. of ferric chloride. 1 mM ATP was also included in all meals. Both sugar and supplemented meal were replaced with fresh meal every 24 hr. After 2-3 days of supplementation, fat-body tissue was collected from 20-25 mosquitoes from both the control and iron supplemented group to check the transcriptional response of selected transcripts.

### *dsRNA* mediated gene knockdown

For *AcTrf1a* knockdown, initially we amplified the single-stranded complementary DNA by applying PCR amplification strategy using dsRNA primers carrying T7 overhang (Supplementary Table 3). The purified PCR product was quantified (Nanodrop 2000 spectrophotometer, Thermo scientific), and validated (agarose gel electrophoresis) using Thermo Scientific Gene JET PCR Purification Kit (Cat #K0701). Purified dscDNA was subjected to double-stranded RNA synthesis using Transcript Aid T7 high-yield transcription kit (Cat# K044, Ambion, USA). After purification, ∼69 nl (∼3ug/ul) of purified dsRNA product was injected into the thorax of newly emerged (1-2 day old) and cold anesthetized mosquitoes using nano-injector facility (Drummond Scientific, CA, USA). Equal number of age matched mosquitoes were also injected with *dsLacZ* (bacterial origin) as control group. 3-4-days post-injection desired samples such as fat-body and ovary were collected from 20-25 mosquitoes, and examined for silencing efficiency by quantitative PCR.

### Mosquito tissue sample collection

Before tissue collection, adult mosquitoes were anesthetized by putting them at 4° C for 4-5 min. Later, placed on to dissecting slide under microscope and various tissue like fat-body, hemocytes, mid gut, salivary gland, ovary, spermatheca and male reproductive organs were collected as described earlier (34). For hemolymph collection, approx. 2-3 μl of anticoagulant consisting of 60% Schneider’s medium, 10% fetal bovine serum, and 30% citrate buffer is injected into thorax. Mosquito belly bulged out, and then a small incision was made in the abdomen using microscopic needle so that transparent hemolymph oozes out. Hemolymph was collected using micropipette and pooled in Trizol. Fat- body was collected by pulling all the abdominal tissues from last two abdominal segments, and afterwards force tapping of abdominal carcass was done so that pale yellow color fat-body oozes. Developmental stages (egg, four larval instars and pupa) were collected independently in Trizol.

### Total RNA isolation and cDNA synthesis

Trizol and alcohol extraction method was used for isolation of total RNA manually from pooled samples, as previously described (35). After quantification by Nanodrop 2000 spectrophotometer (Thermo scientific), 1 μg of total RNA extracted was subjected for synthesis of first strand of cDNA using verso cDNA synthesis kit (Thermo Scientific, USA). Actin was used as a reference gene for quality validation of sscDNA.

### Gene expression analysis

Differential expression of selected transcripts was done using RT-PCR. Primers were designed (https://primer3plus.com/) using either sequenced cDNA or vector base extracted sequences (Supplementary Table 3). SYBR green qPCR master mix and Bio-Rad real time machine was used for relative expression analysis of selected targets under different biological conditions. qPCR protocols remained same for all set of primers with initial denaturation at 95°C for 5 min, 40 cycles of 10 sec at 95°C, 15 sec at 52°C, and 22 sec at 72°C. At the end of each cycle, fluorescence reading was recorded at 72°C. In final steps, PCR at 95°C for 15 sec followed by 55°C for 15 sec and again 95°C for 15 sec were completed before generating melting curve. Three independent biological replicates for each experiment were run using the above protocol. Actin gene was used as an internal control (housekeeping genes) and the relative quantification data were analysed by 2^−ΔΔCt^ method (36). And final relative expression graphs were plotted using origin 8.1 and significance of treatment was determined by paired Student’s *t-*test

### Assessment of ovary development

All experimental groups including control, iron supplemented and silenced were offered blood meal on rabbit. Only fully fed mosquitoes were selected for further experimentation, while partial fed or unfed mosquitoes were discarded. About 72 hr post blood meal, mosquitoes were anesthetized and placed over dissecting slide under binocular microscope for ovary development assessment. Ovaries were dissected in PBS and number of oocytes developed inside the matured ovary were counted manually and compared with control mosquitoes.

### Neutral red staining

Mosquitoes from both control and *AcTrf1a* silenced regime were blood-fed as previously described. Mosquitoes were anesthetized prior to dissection by chilling at 4°C for 3-5 min. At 72 hr of blood meal, ovaries were dissected in 1x PBS and stained with 0.5% neutral red solution in acetate buffer (Sigma–Aldrich, St. Louis, MO). After staining, ovaries were again rinsed in PBS buffer and placed under coverslip. Later these ovaries were visualized under microscope for previtellogenic imaging of ovary.

### Statistical analysis

For statistical analysis, test sample data were compared with the control data set using Origin8.1 software and treatment differences were determined by paired Student’s *t*-test. But wherever required, one-way ANOVA was used for multiple comparisons. The *Mann – Whitney U test* was used to evaluate and analyse the number of developing follicles inside ovary of gravid female mosquitoes. Final *p*-value of less than 0.05 level was considered as significant for both tests. All experiments were conducted thrice for data validation.

## Results

### 1. Molecular characterization of *AcTrf1a*

While analysing RNA-Seq data of the hemocyte, we identified two transcripts, encoding putative transferrin homolog proteins, from the mosquito *An. culicifacies* (under review). An initial BLASTx homology search against non-redundant database indicated that both transcripts encode transferrin domain (Fig. 2a). In this analysis, we observed that only one transcript (2102bp) carries the highest homology to insects (transferrin1) but not the other (2162 bp) which showed unusual clustering with vertebrate’s transferrin proteins (Supplementary Figure 1). Here, we analysed the nature and function of the insect’s transferrin homolog transcript in detail. BLASTn analysis against draft genome database of the mosquito *An. culicifacies*, showed 99% identity to the predicted transcript (ACUA023913-RA), with at least four single nucleotide substitution (Supplementary Table 4). This observation suggested the hemocyte originated transcript is an allelic variant of genome predicted (ACUA023913-RA) transcript, with slight modification of amino acid (aa) sequences (Supplementary Table 4). A comparative physiochemical properties analysis with genome predicted full-length transcript (ACUA023913-RA), indicated that identified *AcTrf1a* is partial in nature, and lack signal peptide sequence (Supplemental Table 2). Multiple-sequence alignment of *AcTrf1a* with its homologs from diverse a class of insects, showed a high degree of sequence conservation among blood feeder mosquitoes (Fig. 2b). The phylogenetic tree obtained at maximum is shown (Fig. 2c). We observed a close relatedness of *AcTrf1a* to blood-feeder insects, by forming a cluster with mosquito transferrin, then non-blood feeder insect clades. (Fig. 2c).

**Fig 2:**
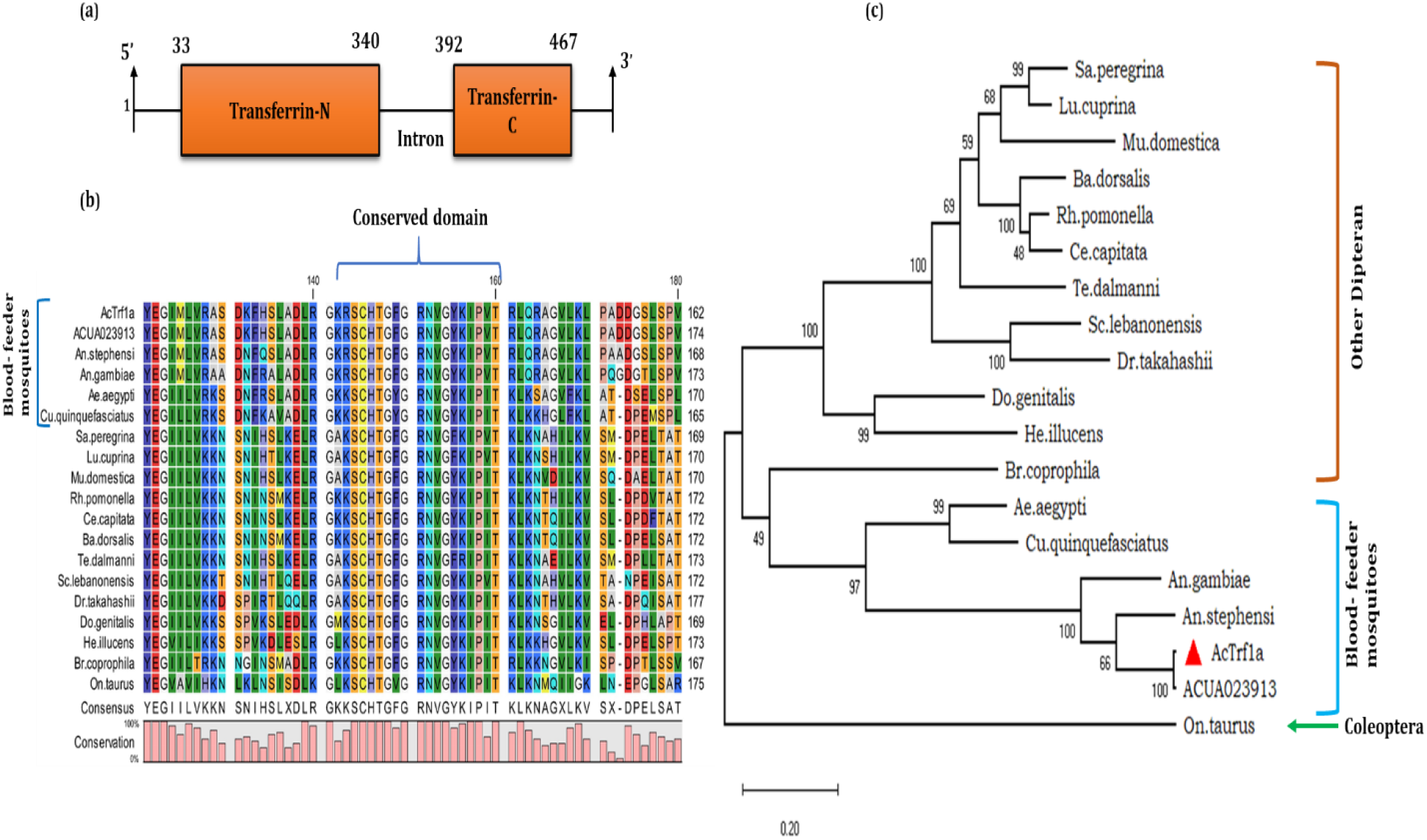
Sequence analysis of *An. culicifacies* hemocytes transcript encoding transferrin1 protein: (a) Pfam (http://pfam.xfam.org) assigned domains annotation of *An. culicifacies* transferrin1 (*AcTrf1a);* (b) The multiple sequence alignment highlights a strong conservation of TRF domain region among the selected insect transferrin homolog proteins; (c) Phylogenetic analysis of the selected transcripts was done using maximum likelihood algorithm upon 1000 times bootstraps in MEGAX software. The percentage of trees in which the associated taxa clustered together is shown next to the branches. The tree is drawn to scale, with branch lengths measured in the number of substitutions per site. *Onthophagus taurus* (coleoptera) transferrin sequence (720 aa) was taken as outgroup for analysis. *AcTrf1a*, allelic form of genome predicted full length transcript ACUA023913-RA, of the mosquito *An. culicifacies*; *An. gambiae* (XP_310734.4); *An. stephensi* (XP_035908482.1); *Ae. aegypti* (AAB87414.1); *Cu. quinquefasciatus* (EDS42355.1); *Sa. peregrina* (Q26643.1); *Lu. cuprina* (XP_023295921.1); *Mu. domestica* (ADU25046.1), *Rh. pomonella* (XP_036318323.1); *Ce. capitate* (XP_004524870.1; *Ba. dorsalis* (AIA24538.1); *Te. dalmanni* (XP_037937085.1); *Sc. lebanonensis* (XP_030379051.1); *Do. genitalis* (AYV99626.1), *He. illucens* (XP_037919193.1); *Br. coprophila* (XP_037051978.1); *Dr. takahashii* (XP_017002431.1); *On. taurus* (XP_022906419.1).

Our domain prediction and annotation analysis indicated that *AcTrf1a* (Fig. 2a), encodes a functional N-terminal lobe (domain), capable of iron-binding. Whereas C-terminal domain is partial/ truncated and most of the known (conserved) iron-binding residues are missing in this domain (Supplementary Table 5). Structural modelling further verifies that the N-terminal transferrin domain carries a regular lobe-like structure (Fig. 3a and Supplementary Table 5) along with cavity and conserved residues necessary for iron-binding. We also noticed that C-terminal domain is much more elongated i.e. less compact, than N-terminal domain, and lack iron-binding site residues and cavity (Fig. 3b and Supplementary Table 5). However, bioinformatic software-based prediction of two putative N-glycosylation sites at asparagine (asn, N) amino acid residue in the C-terminal lobe of transferrin1 at 493 and 500 position of amino acid. (Fig. 3b and Supplementary Table 4), indicated that this newly identified *AcTrf1a*, is an allelic variant of ACUA023913-RA transcript. The secondary structure arrangement of the full protein sequence is shown in Fig. 3c. The modelled *AcTrf1a* structure is composed of 31 helices, and total of 18 cystine residues all are linked by disulphide bonds.

**Fig 3:**
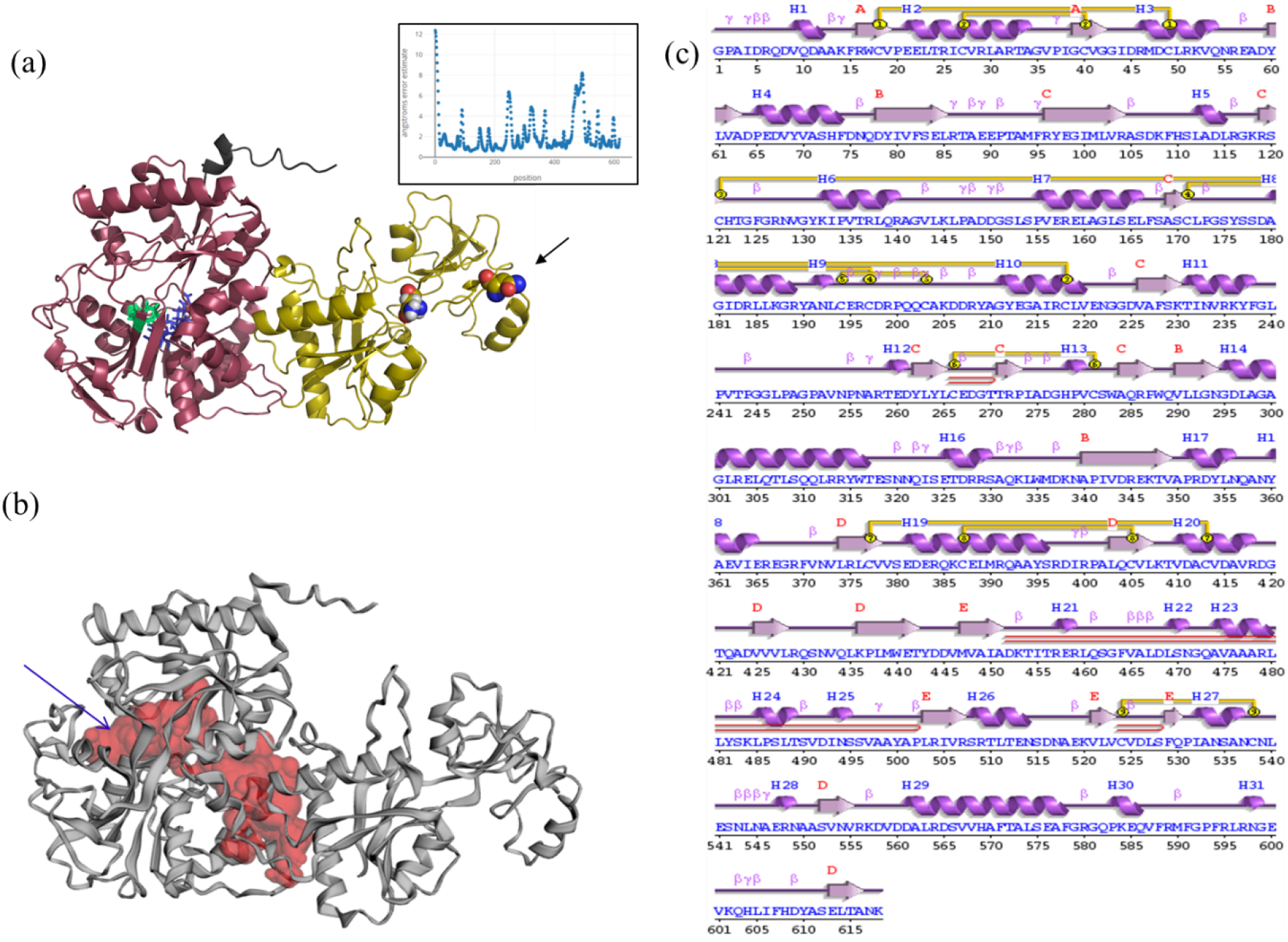
Domain search, localization and structural modeling of *AcTrf1a*. (a) Cartoon representation of *AcTrf1a*, protein model was generated using web-based protein modelling server Robetta (https://robetta.bakerlab.org). In the cartoon, N-terminal domain (N-lobe) of transferrin is shown in dark red, while iron and anion binding residues in green and blue respectively, inset graph is showing the reliability of the protein model across its sequence directly obtained from Robetta (https://robetta.bakerlab.org) server. Further possible glycosylation modification sites were represented in multi-color sphere (asparagine residues) and interestingly both the glycosylation sites were found in C-terminal lobe of *AcTrf1a*, arrow indicates site with superior glycosylation activity. pymol (http://www.pymol.org) was used for visualization purpose; (b) Cavity analysis of *AcTrf1a* was carried out using Castp server (http://sts.bioe.uic.edu/castp/index.html?2cpk), the biggest pocket present in N-lobe encompasses iron-binding site; (c) The output from PDBsum (http://www.ebi.ac.uk/thornton-srv/databases/cgi-bin/pdbsum/GetPage.pl?pdbcode=index.html) runs on sequence of *AcTrf1a* shows secondary structural element present in the modelled protein along with locations of cystine residues.

### 2. Developmental and tissue-specific transcriptional response of *AcTrf1a*

A mosquito developmental stage-specific transcriptional profiling of *AcTrf1a* unveils markedly enriched expression in the eggs and adult female body (*p*<*0.00170*) than other life stages (Fig. 4a) in the *An. culicifacies*. A tissue-specific transcriptional profiling of *AcTrf1*a showed a dominant expression in fat-body (*p*<*0.000213*) and hemocyte (*p*<*0.000524*) (Fig. 4b), then other tissues such as midgut and salivary glands of the 3-4 days old naive adult female mosquitoes. While a sex-specific expression analysis showed that *AcTrf1a* abundantly express in female ovary (*p*<*0.00173*) followed by the male reproductive organ (Fig. 4c). These data suggested that co-expression of *AcTrf1a* in the fat-body and reproductive organ of the naïve adult female mosquitoes may likely have an important role in mosquito reproduction and egg development.

**Fig 4:**
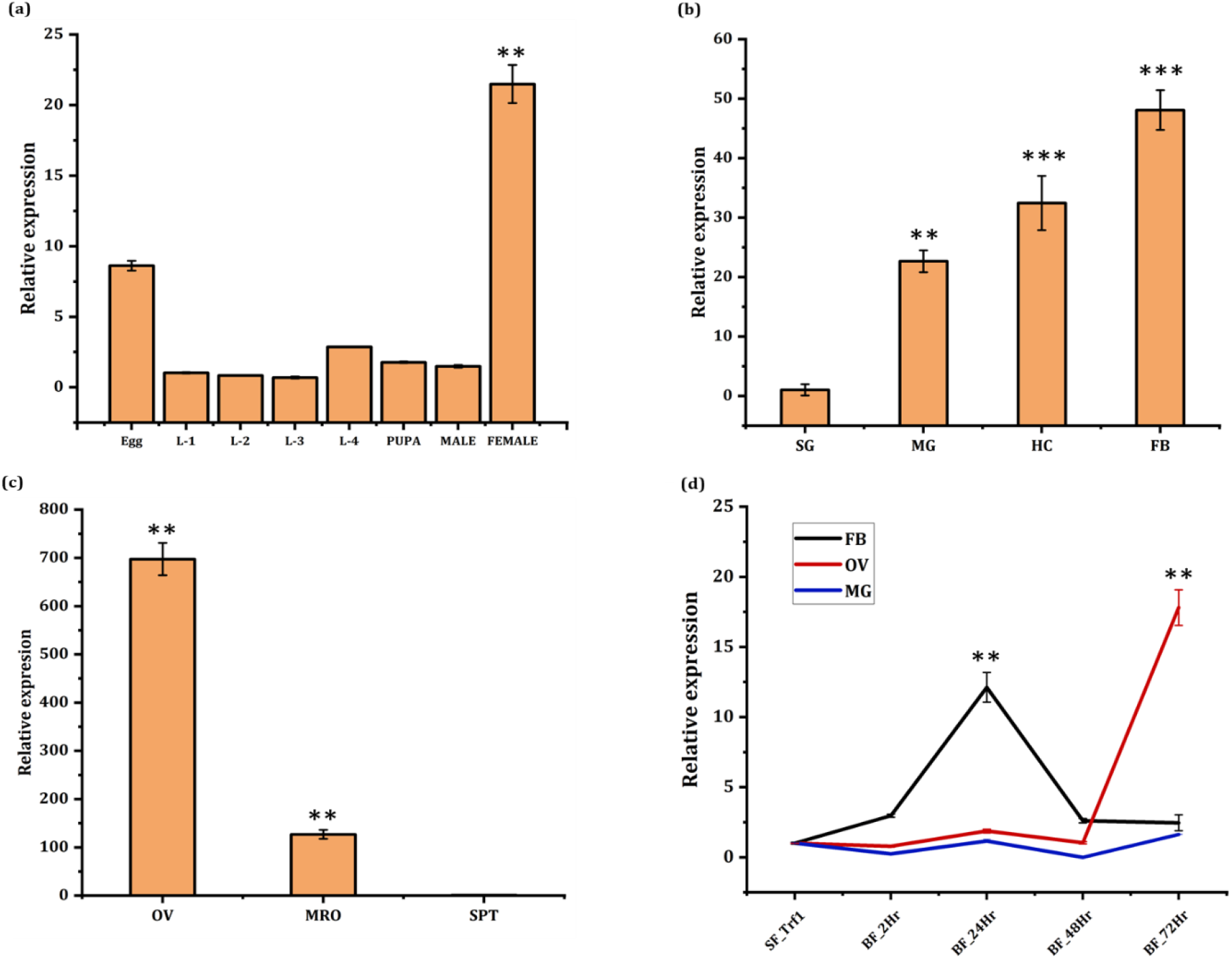
Spatial-Temporal expression profiling of *AcTrf1a* in the mosquito *An. culicifacies*. (a) Relative gene expression during mosquito development, showing an abundance of *AcTrf1a* transcript in the eggs and adult female mosquito (*p*<*0.00170*). First instar larval stage was considered as control for all test sample. L1-4 (larval stages) (n=10, N=3); (b) Expression analysis in the adult female mosquito tissues showing multi-fold upregulation of *AcTrf1a* in fat-body (FB/*p*<*0.000213*), hemocytes (HC/*p*<*0.000524*) and midgut (MG/*p*<*0.00247*) respectively. Data was analysed relative to salivary gland (SG) (n=25, N=3); (c) *AcTrf1a* mRNA levels profiling in the mosquito reproductive tissues shows a prolific expression in ovary (OV/*p*<*0.00173*) and male reproductive tissue (MRO/*p*<*0.00305*). Female spermatheca was set as control for relative analysis (n=20, N=3); (d) Tissue-specific transcriptional response of *AcTrf1a* after blood-feeding: mosquito tissues such as midgut, fat-body and ovary were collected at 2, 24, 48 and 72 hr post blood meal (pbm) and compared with sugar-fed naïve female as a control (n=25, N3). *AcTrf1a* mRNA is minimally expressed in blood-fed midgut, moderate in fat-body (*p*<*0.00226*) post 24 hr of blood meal and highest in 72 hr pbm ovary (*p*<*0.001416*). Three independent biological replicates were performed for statistical analysis, *viz*. *p < 0.05; **p < 0.005; and ***p < 0.0005, using the Student’s *t*-test. (*n*=represents the number of mosquitoes pooled for sample collection; *N*=number of replicates).

### 3. Spatio-temporal expression pattern of *AcTrf1a* following blood-feeding

Host blood meal serve as a rich source of resources for vitellogenesis and follicles maturation in adult female mosquitoes. We explored the possible effect of blood meal, a time-dependent transcriptional response of *AcTrf1a* was monitored in the target tissues such as midgut, fat-body and ovary, ‘*engaged in*’ digestion and nutrient uptake. 3-4-days old adult female mosquitoes were fed on rabbit, and fully engorged females were dissected for the tissues collection at desired time after blood meal. Compared to midgut, a significant modulation in the expression of *AcTrf1a* was observed in the fat-body and ovary. A gradual enrichment in the *AcTrf1a* transcript was observed in fat-body (*p*<*0.00226*) till 24 hr post blood meal, which was restored to basal level within 48 hr of blood-feeding. However, in ovary we observed heightened expression level of *AcTrf1a* post 72 hr (*p*<*0.001416*) of blood meal (Fig. 4d). All together, these data correlate a bi-phasic regulation of transferrin expression of *AcTrf1a* in the fat-body and ovary is essential, possibly to meet the requirement of iron during ovary maturation and egg development, though further investigation are needed.

### 4. Exogenous bacterial challenge differentially regulates *AcTrf1a*

Several studies highlight that insect transferrin serve as an acute-phase protein and play important anti-bacterial immune role (15, 22). Therefore, we also tested whether *AcTrf1a* expression is modulated in response to exogenous bacterial challenge in the naïve mosquito. We injected 2-3 days old adult female mosquito of *An. culicifacies* with heat-killed bacterial suspension of *E. coli* and *B. subtilis* (OD_600_ = 0.9-1). Injections of saline water was considered as control for baseline expression. Post injection, hemocytes were collected from 30-35 mosquitoes at different time interval. We observed that irrespective of the nature of injected bacteria gram-negative or gram-positive (Fig. 5a, b), *AcTrf1a* transcript expressed highly within two hrs of bacterial challenge was recorded. Together, these observations further strengthen the hypothesis that *AcTrf1a* have antibacterial role in the mosquito *An. culicifacies*.

**Fig 5:**
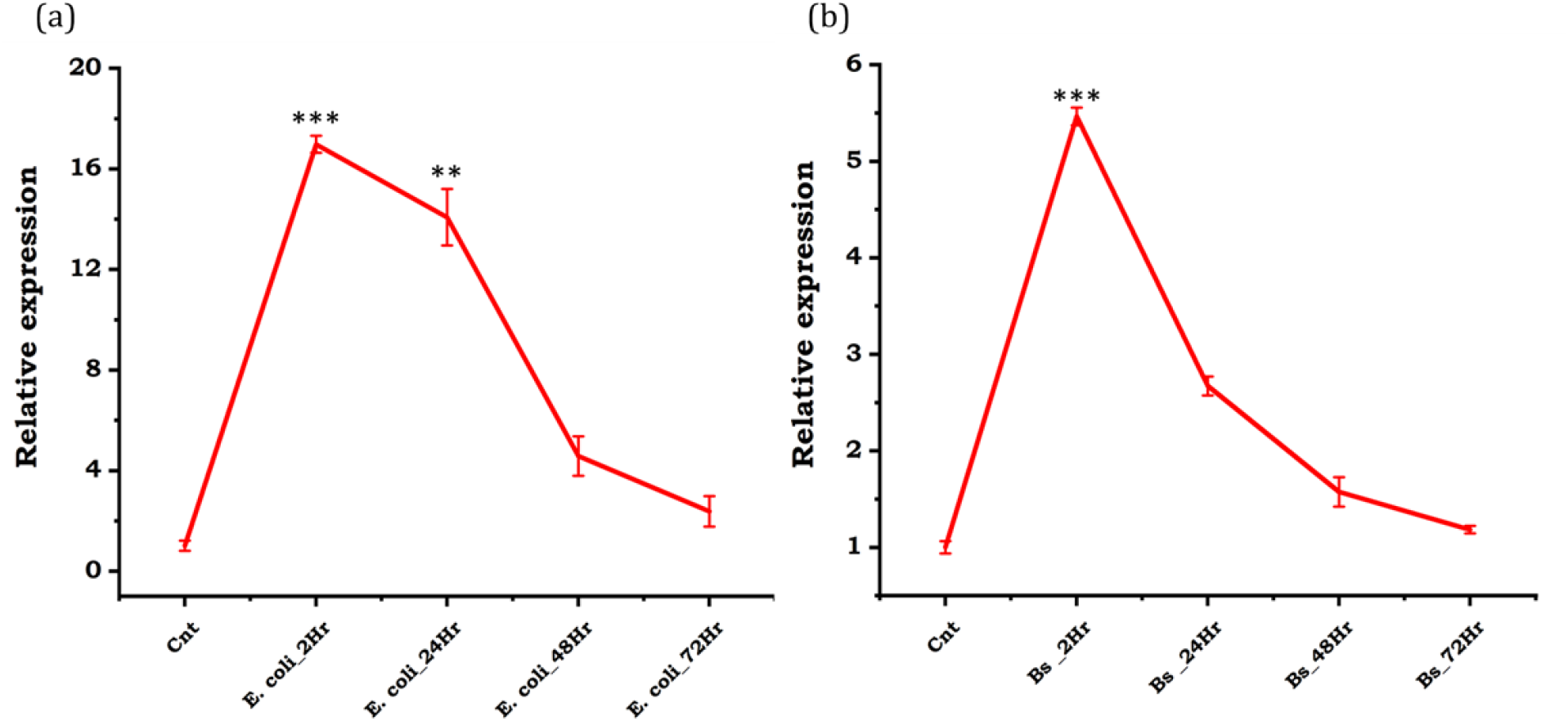
Transcriptional response of *AcTrf1a* against bacterial challenge. Adult female mosquito of 3-4 days old was injected either with 69 nl of heat killed bacterial suspension in PBS or PBS alone as a control in mosquito thorax (n=30-35, N3). After recovery, time dependent series, hemocyte samples were collected post (2, 24, 48 & 72 hr) bacterial challenge. (a) Relative transcriptional profiling against *E. coli* (*p*<*0.000359*) and; (b) *Bacillus subtilis* (*p*<*0.000156*) shows the strong antibacterial nature of *AcTrf1a*. Statistical analysis was done using student’s *t*- test, *viz* *p < 0.05; **p < 0.005; and ***p < 0.0005, (*n*=represents the number of mosquitoes pooled for sample collection; *N*= number of replicates); pbc, post bacterial challenge; PBS, phosphate saline buffer; Bs, *Bacillus subtilis*.

### 5. Effect of iron supplement on *AcTrf1a* expression and oocyte load

Next, to test the possible role of *AcTrf1a* in the reproductive physiology, first we evaluated whether supplementation of non-heme iron modulate the mosquito transferrin’s expression. To ensure an optimal impact, initially we offered two independent regimes of different serial dilution of ferric chloride i.e., 1 μM, 5 μM, 10 μM, and 0.1 mM, 0.5 Mm, 1 mM in 10% sugar solution female mosquito just after emergence *via* oral feeding. After 48-72 hr of iron feeding, fat-body tissues were collected and the transcription profile of *AcTrf1a* was monitored through Real-Time PCR assay. Compared to low concentration, which showed a mild impact (Supplementary Fig. 2), a significant upregulation was noticed in the naïve mosquitoes, when fed with a concentration of 1 mM FeCl_3_ (Fig. 6a).

**Fig 6:**
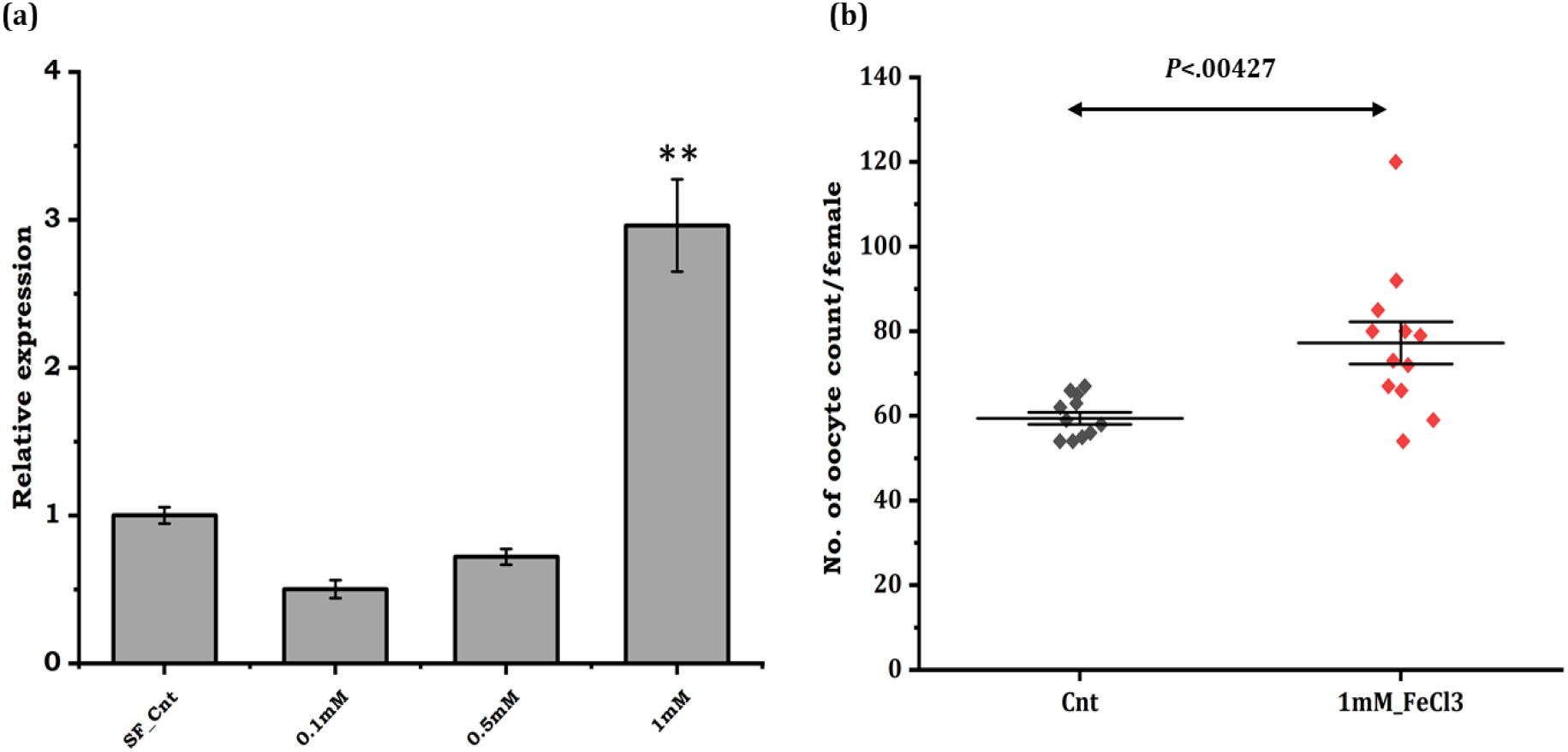
mRNA level augmentation of *AcTrf1a* transcripts on iron supplemented diet and effect on the mosquito fecundity. Female mosquitoes were allowed to feed on different iron concentration prepared in sugar solution and mRNA level of target transcripts was measured in the fat-body. (a) Comparative transcriptional profiling of *AcTrf1a* in fat-body tissue collected from control and treatment group. An enriched expression (*p*<*0.00508*) was observed at 1 mM conc. of iron supplemented diet. All experiments were performed in triplicate for statistical analysis; using student’s *t*-test; (n=20-25, N=3) (b) Dot plot showing that iron supplemented diet during the early stage of development just after eclosion leads to enrichment of oocytes (*p*<*0.00427*) maturing inside the female ovary. Both the control (SF) and test group (SF+ FeCl_3_) were allowed to feed on rabbit blood and subsequently kept for ovary assessment for oocyte count (n=10, N=3). Data represents mean of three biological replicates and significance was analysed using Mann- Whitney *U* test. (*n*=represents the number of mosquitoes pooled for sample collection; *N*= number of replicates). *p<0.05; **p<0.005 and ***p<0.0005.

With this optimization, we investigate whether this extra iron supplementation (1 mM FeCl_3_) influences mosquito reproductive physiology such as the number of developing oocytes inside the ovary. To perform this assay, mosquitoes were kept under two nutritional status: 1) 10% sugar solution only or 2) 10% sugar supplemented with ferric chloride solution (1 mM). Post supplementation, both groups of mosquitoes were offered rabbit blood for successful completion of first gonotrophic cycle. After 72 hr of blood meal, ten mosquitoes from each control and test group were dissected and examined for ovary development. Mosquitoes fed with 10% sugar alone had an average 60 mature oocytes per female. While mosquito kept on iron supplemented diet, yielded an average of 75 oocytes/female mosquito (*p*<*0.00508)* (Fig. 6b). This data highlighted that *AcTrf1a* may have an important role in the mosquito reproductive physiology, possibly by altering pre-vitellogenic nutritional iron transport and status of resting stage.

### 6. Transferrin knockdown reduces oocyte count in ovary

To further evaluate and support the above hypothesis, we performed dsRNA mediated gene knockdown/silencing experiment. For this, 2-day old mosquitoes were injected with double-stranded RNA of *AcTrf1a* and mosquitoes were kept under optimal insectary conditions for better recovery. Fat-body and ovary tissues were collected from 20-25 mosquito three-day post silencing to check the transcript mRNA level. Compared to *dsLacZ* injected control mosquito group, we observed at least 70% reduction in the transcript of *AcTrf1a* expression in the fat-body (*p*<*0.000320*) and 50% in the ovary (*p*<*0.00174*) of knockdown mosquitoes (Fig. 7a). Both control and silenced group were offered blood meal and after 72hr of blood meal, individual mosquito (n=10) was dissected in PBS and total number of mature oocytes were manually counted under microscope. Our initial phase contrast microscopic analysis showed significant difference in maturing follicles after silencing (Supplementary figure 3). We observed that in contrast to control mosquito which have 63 (average) oocyte per mosquito, silenced mosquitoes showed only 40 (average) oocytes per mosquito (Fig. 7b). Our data suggested that *AcTrf1a* has direct influence on the mosquito reproductive physiology.

**Fig. 7:**
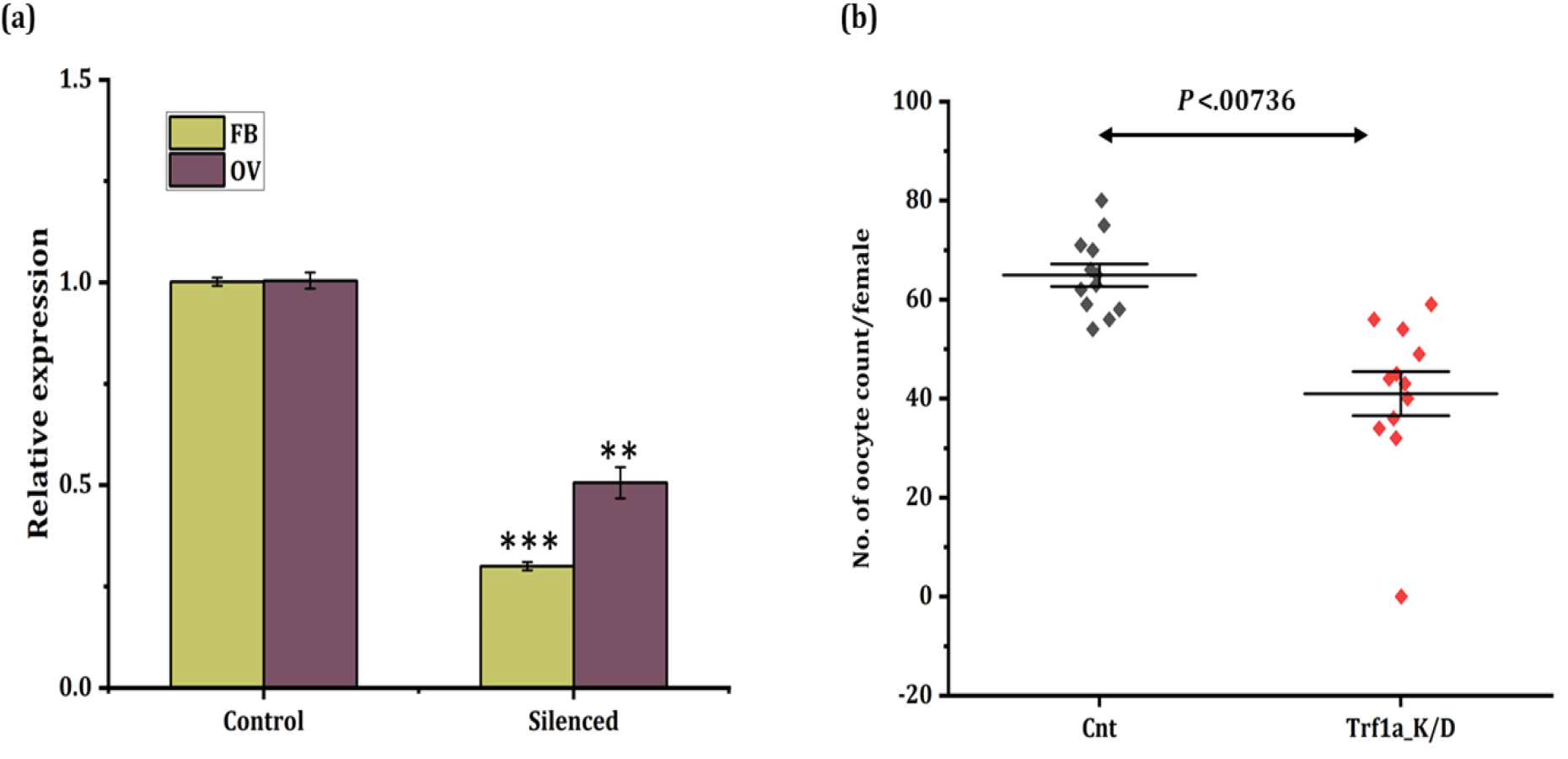
*AcTrf1a* silencing effect on mosquito oocyte development. (a) Real time-PCR based knockdown validation of *AcTrf1a* in fat-body (*p*<*0.000320*) and ovary (*p*<*0.00174*) tissue after *dsrAcTrf1a* injection to newly emerged mosquitoes in comparison to same age group *dsrLacZ* injected (n=25, N=3); (b) Dot plot showing the comparative oocyte count reduction (*p*<*0.00736*) post *AcTrf1a* knockdown in female ovary tissue compared to control mosquito group. Post dsRNA injection mosquitoes were fed on blood meal and ovary assessment was done, (n=10, *N*=3). Statistical significance *p<0.05; **p<0.005 and ***p<0.0005 was calculated using student’s t-test and Mann- Whitney *U* test. (*n*=represents the number of mosquitoes pooled for sample collection; *N*= number of replicates). FB, fat-body; OV, ovary.

## Discussion

Transferrin family is a large group of protein found in both vertebrates as well as invertebrates, including insects, and play an important role in the iron transportation and metabolism (37–40). The majority of family members have two lobes for iron-binding which evolved *via* gene duplication event during the course of evolution (12, 41). Transferrin protein is pleiotropic in insects, involved in iron homeostasis, immunity, vitellogenic protein, antibacterial agent, and sequestered uptake in ovarian follicles (15, 17, 42–44).

Although, several studies highlight antibacterial role of transferrin in the adult female mosquitoes, that feed on iron-rich blood, but its role in ovarian development and egg maturation, remains poorly understood. Here, we have identified and characterized an iron-binding transferrin allelic variant *(AcTrf1a)* from the hemocyte transcriptome of Indian malarial vector *An. culicifacies*. Through comprehensive spatial/temporal expressional profiling, and functional knockout studies, for the first time we demonstrate that mosquito transferrin significantly influence the reproductive physiology of the mosquito *An. culicifacies*. The initial finding of transferrin transcripts from the hemocyte transcriptome corroborates with previous observation that the transferrin is expressed dominantly in the fat-body, and haemolymph (13, 16). Unlike, vertebrates and insects, the blood-feeding mosquito transferrin lacks N-Glycosylation site (15). But prediction of N-Glycosylation site in the hemocyte originated transferrin (*AcTrf1a)*, indicates that this is an allelic variant of ACUA023913-RA (genome predicted transcript), which shares 99.36% identity at amino-acid level (Supplementary Table 6). Recent structural studies of insect transferrin show that iron-binding mechanism of insect transferrin (*Trf1*) is completely different compared to another known vertebrate transferrin. Insects transferrin (*Manduca sexta*) iron-binding site is localized at N-terminus and coordinated by two tyrosine ligands, and two CO_3_ ^2-^ anions, contrary to vertebrate transferrin where both N- and C-lobe participate in iron-binding (45). Furthermore, in vertebrate transferrin iron-binding site is constituted of two tyrosines, one aspartate, one histidine, and one carbonate anion (46, 47). A dominant expression of *AcTrf1a* in the egg than other developmental stages such as larva, pupae, and in 3-4 days old adult female than age matched male mosquitoes, together indicates that *AcTrf1a* may play key role in the egg development of this mosquito species. A similar expression pattern for other iron transport protein i.e., ferritin has also been observed in *Aedes* (48). Compared to fat-body/hemocyte, an observation of a multifold enriched expression in the reproductive organs of both sexes, further strengthen the hypothesis that *AcTrf1a* may likely have potential role in the transportation, and optimal iron supply maintenance in the sexually maturing reproductive organs.

In hematophagous insects, intake of iron rich blood meal is crucial for oocyte development, ovary maturation and successful completion of gonotrophic cycle (5, 49). Although our 3D model and structure prediction analysis confer iron and anion binding site/pocket at N-terminal of putative *AcTrf1a*. However, the lack of vertebrate transferrin receptor homolog in insects/mosquitoes, it remains unresolved that how loading and unloading of iron is achieved at midgut and ovary surfaces, respectively (8, 50). A recent study by Farkas et.al 2018 suggests the involvement of basally derived endosomes in iron import to the target cells is important in the insect *D. melanogaster*, and proposes that basal endocytosis is followed by vacuole acidification, a process leading to the activation of ferrireductase for iron release and metabolic utilization (51). Several studies highlight that transferrin play an antibacterial immune role against microbial infection in insects (52–54). Constitutive expression in the fat-body and haemolymph, and a significant upregulation upon bacterial challenge, together indicates that the mosquito transferrin *AcTrf1a* may also have antibacterial role, similar to vertebrate lactotransferrin (55).

Next, we noticed that *AcTrf1a* transcript is upregulated after 24 hr of blood meal in the fat-body, which may likely facilitate to meet the iron supply for ovarian follicle development during the active vitellogenic process. While endogenous expression enrichment of *Actrf1a* in the ovary after 72 hr of blood-feeding is necessary for the egg development in the matured oocytes. Possibly, this is achieved by uptake of iron from hemolymph as well as synthesizing endogenously in ovary, though further studies needed to validate these propositions. Kumara et. al. also reported the sequestered uptake of transferrin by developing oocytes in flesh fly (17). Next, we tested whether exogenous iron supply through oral supplementation of non-heme iron alters the *AcTrf1a* expression in the fat-body and midgut of the naïve adult female mosquito. Here, we observed that the expression remains suppressed until an optimal concentration (1 mM) triggers the upregulation of *AcTrf1a* in the fat-body of the naïve mosquito. Corroborating with the previous studies these findings suggest that *AcTrf1a* may also have role as an antioxidant molecule in the blood-feeding insects (22, 56). We hypothesized that this iron supplementation, and natural blood-feeding together may lead to a synergistic effect on the transcript expression modulation, accelerating iron transportation and ultimately enhancing the reproductive potential.

Supporting the above hypothesis, our data highlighted that oral supplementation of non-heme iron not only upregulate the transcriptional level, but also augment the oocyte load in the ovary of gravid adult female mosquitoes. Earlier studies in *Aedes aegypti*, shows that exogenous supply boosts the transferrin expression, however, its functional correlation to ovarian development remain uncertain. Our functional knockdown experiment showed that there is a significant reduction in oocyte load in the developing ovaries. Taken together, our data suggests that transferrin performs two major functions of iron transport and oxidative stress management, and thus may serve as an essential molecule to meet optimal iron supply during oogenesis process.

## Conclusion

In summary, our results are consistent with previous studies in other mosquitoes and insects suggesting transferrin as a multifunctional protein involved in iron transportation, antibacterial immune protein, and vitellogenesis. We speculated that transferrin may have a physiological role in trade-off of resources between immunity and reproduction. Possibly this mechanism may facilitate an optimal storage and transportation of iron from the midgut/fat-body to the ovaries. Here, we propose that hemocyte originated allelic variant of transferrin plays a pivotal role in mosquito fecundity and survival.

## Acknowledgement

We thank all staff members of insectary for mosquito rearing and Kunwarjeet Singh for technical support in laboratory. We all also thankful to animal house staff members for animal (rabbit) support, whenever required. Finally, we thank to NGB Diagnostics Pvt. Ltd., Delhi for generating sequencing data.

## Authors contribution statement

JR, NS and RD conceived and designed the experiments; CC, TTD, SK, PS, ST, SC helped performing experiments, drafting and editing MS; NS, RD, KCP Contributed reagents/materials/analysis tools, helped in writing, editing, reviewing and finalizing MS. All authors read and approved the final manuscript.

## References

1. B Dunkov; T Georgieva. Insect iron binding proteins: Insights from the genomes. Insect Biochem Mol Biol 36, 300–309 (2006)

2. M Muñoz; I Villar; JA García-Erce. An update on iron physiology. World J Gastroenterol 15, 4617–4626 (2009)

3. AK Hernández-Gallardo; F Missirlis. Loss of ferritin in developing wing cells: Apoptosis and ferroptosis coincide. PLoS Genet 16, 2–7 (2020)

4. C Rivera-Pérez; ME Clifton; FG Noriega. How micronutrients influence the physiology of mosquitoes, (2017)

5. R Jason Pitts. A blood-free protein meal supporting oogenesis in the Asian tiger mosquito, Aedes albopictus (Skuse). J Insect Physiol (2014)

6. GD Id; O Banmeke; YZ Id; S Huang; M Hamilton; Y Ping; C Id. Transferrin-mediated iron sequestration suggests a novel therapeutic strategy for controlling Nosema disease in the honey bee, Apis mellifera. 1–30 (2021)

7. DB Kell. Iron behaving badly: Inappropriate iron chelation as a major contributor to the aetiology of vascular and other progressive inflammatory and degenerative diseases. BMC Med Genomics 2, 1–79 (2009)

8. X Tang; B Zhou. Iron homeostasis in insects: Insights from Drosophila studies. IUBMB Life 65, 863–872 (2013)

9. S Recalcati; E Gammella; P Buratti; G Cairo. Molecular regulation of cellular iron balance, (2017)

10. SP Young; P Aisen. Transferrin receptors and the uptake and release of iron by isolated hepatocytes. Hepatology 1, 114–119 (1981)

11. EL Mackenzie; K Iwasaki; Y Tsuji. Intracellular iron transport and storage: From molecular mechanisms to health implications, (2008)

12. LA Lambert; H Perri; TJ Meehan. Evolution of duplications in the transferrin family of proteins. Comp Biochem Physiol - B Biochem Mol Biol 140, 11–25 (2005)

13. G Xiao; ZH Liu; M Zhao; HL Wang; B Zhou. Transferrin 1 Functions in Iron Trafficking and Genetically Interacts with Ferritin in Drosophila melanogaster. Cell Rep 26, 748–758.e5 (2019)

14. LA Lambert; H Perri; PJ Halbrooks; AB Mason. Evolution of the transferrin family: Conservation of residues associated with iron and anion binding, (2005)

15. DL Geiser; JJ Winzerling. Insect transferrins: Multifunctional proteins. Biochim Biophys Acta - Gen Subj 1820, 437–451 (2012)

16. N Harizanova; M Tchorbadjieva; P Ivanova; S Dimov; K Ralchev. Developmental and organ-specific expression of transferrin in drosophila melanogaster. Biotechnol Biotechnol Equip 18, 118–121 (2004)

17. T Kurama; S Kurata; S Natori. Molecular Characterization of an Insect Transferrin and its Selective Incorporation into Eggs During Oogenesis. Eur J Biochem 228, 229–235 (1995)

18. M Hirai; D Watanabe; Y Chinzei. A juvenile hormone-repressible transferrin-like protein from the bean bug, Riptortus clavatus: CDNA sequence analysis and protein identification during diapause and vitellogenesis. Arch Insect Biochem Physiol (2000)

19. JB Benoit; GM Attardo; V Michalkova; TB Krause; J Bohova; Q Zhang; AA Baumann; PO Mireji; P Takáč; DL Denlinger; JM Ribeiro; S Aksoy. A Novel Highly Divergent Protein Family Identified from a Viviparous Insect by RNA-seq Analysis: A Potential Target for Tsetse Fly-Specific Abortifacients. PLoS Genet 10, 6–10 (2014)

20. JJ Weber; MR Kanost; MJ Gorman. Iron binding and release properties of transferrin-1 from Drosophila melanogaster and Manduca sexta: Implications for insect iron homeostasis. Insect Biochem Mol Biol 125, 103438 (2020)

21. JH Law. Insects, oxygen, and iron. Biochem Biophys Res Commun (2002)

22. T Yoshiga; VP Hernandez; AM Fallon; JH Law. Mosquito transferrin, an acute-phase protein that is up-regulated upon infection. Proc Natl Acad Sci U S A 94, 12337–12342 (1997)

23. L Ling; AS Raikhel. Cross-talk of insulin-like peptides, juvenile hormone, and 20-hydroxyecdysone in regulation of metabolism in the mosquito Aedes aegypti. Proc Natl Acad Sci (2021)

24. AN Clements. The biology of mosquitoes. (2011)

25. X Tang; B Zhou. Ferritin is the key to dietary iron absorption and tissue iron detoxification in Drosophila melanogaster. FASEB J (2013)

26. J Emerit; C Beaumont; F Trivin. Iron metabolism, free radicals, and oxidative injury, (2001)

27. M Wessling-Resnick. Iron homeostasis and the inflammatory response, (2010)

28. N Harizanova; T Georgieva; BC Dunkov; T Yoshiga; JH Law. Aedes aegypti transferrin. Gene structure, expression pattern, and regulation. Insect Mol Biol 14, 79–88 (2005)

29. G Zhou; LS Velasquez; DL Geiser; JJ Mayo; JJ Winzerling. Differential regulation of transferrin 1 and 2 in Aedes aegypti. Insect Biochem Mol Biol 39, 234–244 (2009)

30. KP Paily; BA Kumar; K Balaraman. Transferrin in the mosquito, Culex quinquefasciatus Say (Diptera: Culicidae), up-regulated upon infection and development of the filarial parasite, Wuchereria bancrofti (Cobbold) (Spirurida: Onchocercidae). Parasitol Res (2007)

31. W Tan; X Wang; P Cheng; L Liu; H Wang; M Gong; X Quan; H Gao; C Zhu. Cloning and overexpression of transferrin gene from cypermethrin-resistant Culex pipiens pallens. Parasitol Res (2012)

32. V Dev; VP Sharma. The Dominant Mosquito Vectors of Human Malaria in India. In: Anopheles mosquitoes - New insights into malaria vectors (2013)

33. S Kumar; G Stecher; M Li; C Knyaz; K Tamura. MEGA X: Molecular evolutionary genetics analysis across computing platforms. Mol Biol Evol (2018)

34. S Kumari; C Chauhan; S Tevatiya; D Singla; T Das De; P Sharma; T Thomas; J Rani; D Savargaonkar; KC Pandey; V Pande; R Dixit. Current Research in Immunology Genetic changes of Plasmodium vivax tempers host tissue-speci fi c responses in Anopheles stephensi. Curr Res Immunol 2, 12–22 (2021)

35. T Das De; P Sharma; T Thomas; D Singla; S Tevatiya; S Kumari; C Chauhan; J Rani; V Srivastava; R Kaur; KC Pandey; R Dixit. Interorgan molecular communication strategies of “Local” and “Systemic” innate immune responses in mosquito Anopheles stephensi. Front Immunol (2018)

36. KJ Livak; TD Schmittgen. Analysis of relative gene expression data using real-time quantitative PCR and the 2-ΔΔCT method. Methods 25, 402–408 (2001)

37. H Kawabata. Transferrin and transferrin receptors update, (2019)

38. L Bai; M Qiao; R Zheng; C Deng; S Mei; W Chen. Phylogenomic analysis of transferrin family from animals and plants. Comp Biochem Physiol - Part D Genomics Proteomics (2016)

39. K Gkouvatsos; G Papanikolaou; K Pantopoulos. Regulation of iron transport and the role of transferrin, (2012)

40. JL Beard. Iron biology in immune function, muscle metabolism and neuronal functioning. In: Journal of Nutrition (2001)

41. H Mohd-Padil; A Mohd-Adnan; T Gabaldón. Phylogenetic analyses uncover a novel clade of transferrin in nonmammalian vertebrates. Mol Biol Evol (2013)

42. R Kucharski; R Maleszka. Transcriptional profiling reveals multifunctional roles for transferrin in the honeybee, Apis mellifera. J Insect Sci 3 (2003)

43. DG Najera; NT Dittmer; JJ Weber; MR Kanost; MJ Gorman. Phylogenetic and sequence analyses of insect transferrins suggest that only transferrin 1 has a role in iron homeostasis. Insect Sci (2020)

44. C Sangokoya; JF Doss; JT Chi. Iron-Responsive miR-485-3p Regulates Cellular Iron Homeostasis by Targeting Ferroportin. PLoS Genet 9, 1–11 (2013)

45. JJ Weber; MM Kashipathy; KP Battaile; E Go; H Desaire; MR Kanost; S Lovell; MJ Gorman. Structural insight into the novel iron-coordination and domain interactions of transferrin-1 from a model insect, Manduca sexta. Protein Sci (2021)

46. K Mizutani; M Toyoda; B Mikami. X-ray structures of transferrins and related proteins, (2012)

47. H Sun; H Li; PJ Sadler. Transferrin as a metal ion mediator. Chem Rev (1999)

48. DL Geiser; TN Thai; MB Love; JJ Winzerling. Iron and ferritin deposition in the ovarian tissues of the yellow fever mosquito (Diptera: Culicidae). J Insect Sci 19 (2019)

49. J Marques; JCR Cardoso; RC Felix; RAG Santana; M das GB Guerra; D Power; H Silveira. Fresh-blood-free diet for rearing malaria mosquito vectors. Sci Rep (2018)

50. K Mandilaras; T Pathmanathan; F Missirlis. Iron absorption in Drosophila melanogaster, (2013)

51. R Farkaš; D Beňová-Liszeková; L Mentelová; M Beňo; K Babišová; L Trusinová-Pečeňová; O Raška; BA Chase; I Raška. Endosomal vacuoles of the prepupal salivary glands of Drosophila play an essential role in the metabolic reallocation of iron. Dev Growth Differ (2018)

52. LM Brummett; MR Kanost; MJ Gorman. The immune properties of Manduca sexta transferrin. Insect Biochem Mol Biol 81, 1–9 (2017)

53. KS Lee; BY Kim; HJ Kim; SJ Seo; HJ Yoon; YS Choi; I Kim; YS Han; YH Je; SM Lee; DH Kim; HD Sohn; BR Jin. Transferrin inhibits stress-induced apoptosis in a beetle. Free Radic Biol Med (2006)

54. EY Yun; JK Lee; OY Kwon; JS Hwang; I Kim; SW Kang; WJ Lee; JL Ding; KH You; TW Goo. Bombyx mori transferrin: Genomic structure, expression and antimicrobial activity of recombinant protein. Dev Comp Immunol (2009)

55. D Legrand. Overview of Lactoferrin as a Natural Immune Modulator. J Pediatr (2016)

56. SR Whiten; H Eggleston; ZN Adelman. Ironing out the details: Exploring the role of iron and heme in blood-sucking arthropods, (2018)

